# Caveolae promote successful abscission by controlling intercellular bridge tension during cytokinesis

**DOI:** 10.1101/2022.01.04.474800

**Authors:** Virginia Andrade, Jian Bai, Neetu Gupta-Rossi, Ana Jimenez, Cédric Delevoye, Christophe Lamaze, Arnaud Echard

## Abstract

During cytokinesis, the intercellular bridge (ICB) connecting the daughter cells experiences pulling forces, which delay abscission by preventing the assembly of the ESCRT scission machinery. Abscission is thus triggered by tension release, but how ICB tension is controlled is unknown. Here, we report that caveolae, which are known to control membrane tension upon mechanical stress in interphase cells, are located at the midbody, at the abscission site and at the ICB/cell interface in dividing cells. Functionally, the loss of caveolae delays ESCRT-III recruitment during cytokinesis and impairs abscission. This is the consequence of a 2-fold increase of ICB tension measured by laser ablation, associated with a local increase in myosin II activity at the ICB/cell interface. We thus propose that caveolae buffer membrane tension and limit contractibility at the ICB to promote ESCRT-III assembly and cytokinetic abscission. Altogether, this work reveals an unexpected connection between caveolae and the ESCRT machinery and the first role of caveolae in cell division.

**TEASER:** Caveolae limit the tension in the intercellular bridge during cytokinesis to enable ESCRT-III assembly and successful abscission.

## INTRODUCTION

Cytokinesis is the last step of cell division and physically separates the daughter cells. In animal cells, cytokinesis is driven by local and highly coordinated polymerization/depolymerization of different types of cytoskeleton and filaments (*1, 2*). First, the ingression of the cleavage furrow depends on the contraction of an acto- myosin ring at the cell equator. Then, the intercellular bridge (ICB) connecting the daughter cells is stabilized by septins. Finally, the midbody at the center of the ICB recruits the abscission machinery that pinches the membrane on one side of the midbody at the future abscission site (**Fig. S1A**) (*1, 2*). This last step requires the local disassembly of microtubules and actin filaments (*2℃9*), as well as the timely polymerization of ESCRT-III filaments at the abscission site (*10–16*). The ESCRT-III-driven constriction of the plasma membrane eventually leads to the scission of the ICB (*2, 12*).

What triggers ESCRT-III fliament assembly at the right place and time, first at the midbody then at the abscission site remains poorly understood. While several biochemical components, such as CEP55, MKLP1, ALIX, TSG101, Syndecan-4, are key for the proper recruitment, localization and polymerization of the ESCRT-III machinery (*2, 12*), it has been shown that mechanical inputs play a critical role as well. Indeed, after furrow ingression, the daughter cells exert nano-Newton (nN)-range pulling forces on the bridge, which is thus under tension (*17*). Rather than favoring abscission, this, on the contrary, delays ESCRT-III recruitment at the abscission site (*17*). It has thus been proposed that abscission requires a tension release in the ICB (*17*), but how this is controlled remains elusive. Understanding the regulation of the intercellular forces in the ICB should thus reveal one of the most upstream events that initiates abscission.

Here, we identified caveolae as a key regulator that limits ICB tension to promote abscission. Caveolae are characteristic cup-shaped 60-80 nm invaginations of the plasma membrane, discovered in the 1950’s (*18–22*). The formation of caveolae requires two types of structural components: Caveolins, which are partially embedded in the plasma membrane, and Cavins that form a coat around caveolae (*20, 21*). In addition, accessory proteins such as EHD2 and PACSINS regulate caveolae dynamics (*20℃22*). In vertebrates, caveolae are particularly abundant in cells that experience chronical mechanical constraints, such as muscle cells, endothelial cells and adipocytes (*20, 21*). Functionally, caveolae define nanometer scale lipid compartments at the plasma membrane and have been involved in the regulation of many signaling events (*23*). Over the past decade, it also emerged that caveolae constitute a membrane reservoir that plays a critical role in buffering membrane tension in non-dividing cells experiencing mechanical stress, through their ability to flatten out (*24–27*). This is particularly important to maintain tissue integrity in adult tissues, during development and in pathological situations such as cancers (*28–33*).

Cells experience dramatic forces and shape changes during cell division, but little is known about caveolae during mitosis or cytokinesis. At the onset of mitosis, before chromosome segregation, caveolin-1 is enriched at cortical regions and guides spindle orientation (*34*). Later, in metaphase, caveolin-1 redistribute from the plasma membrane to internal compartments, accompanied with fission of caveolae from the plasma membrane (*35*). Conversely, caveolin-1 reappears at the membrane during telophase in HeLa cells (*35*). Furthermore, it is known for long that caveolin-1 concentrate at the cleavage furrow in MDCK cells and in early zebrafish embryos (*36, 37*). However, the significance of this dynamic redistribution of caveolin-1 during mitotic exit, and whether caveolae have a role in cell division remains unknown.

Using a variety of approaches, from endogenous immunostaining, high resolution spinning disk confocal microscopy, electron microscopy, CRISPR-Cas9 KO cells to laser ablation for measuring tension, we report that 1) *bona fide* caveolae are present at the midbody as well as at the sides of the ICB after furrow ingression, 2) caveolae regulate acto-myosin II-dependent contractility at the ICB and ICB mechanical tension, and 3) by limiting ICB tension, caveolae promote both ESCRT-III assembly and abscission. This work thus reveals the first function of caveolae in cell division, whereby caveolae buffer tension in the ICB, a physical parameter that is critical for successful cytokinesis.

## RESULTS

### Caveolae are present at the midbody and at the ICB/cell interface

In many cell types, abscission occurs on both sides of the midbody and releases a midbody remnant (MBR) (*17, 38℃42*) (**Fig. S1A**). We recently developed a new method for isolating pure, intact MBRs from HeLa cells and provided the quantitative proteome of this organelle that we termed the *Flemmingsome* (*43*). Data analysis led to the intriguing discovery that several proteins constitutive of caveolae were either present (caveolin-1, PACSIN2, Cavin3 and EHD2) or enriched (Cavin1/PTRF and PACSIN3, >3.5 fold and >3 fold compared to control fractions, respectively) in the *Flemmingsome* (**Fig. S1B**). In this study, we chose to focus on Cavin1 and caveolin-1 (Cav1), as both are essential structural components of caveolae (*20, 21*).

Immunostaining of purified MBRs from HeLa cells expressing the midbody marker GFP-MKLP1 showed endogenous Cavin1 and Cav1 localization on dotty structures (**Fig. 1A and S1C**), consistent with the proteomic data. During cytokinesis, we also observed an accumulation of similar structures containing Cavin1 (**Fig. 1B**) and Cav1 (**Fig. S1D**) at the cytokinetic furrow when ingressed, as previously reported for Cav1 in other cell types (*36, 37*). In addition, we found that both proteins localized to the midbody in 80-90% of both early and late ICBs (characterized by thick and thin tubulin staining, respectively (*39, 43*)), which had not been previously reported thus far (**Fig. 1B and S1D**). The presence of Cavin1 dots suggested the existence of caveolae at the midbody, since Cavin1 at the plasma membrane is only present on fully budded caveolae at the plasma membrane (*22*). To determine whether *bona fide* caveolae were present at the midbody, we turned to electron microscopy (EM). Using scanning EM (SEM), we observed small open structures at the midbody ring that likely correspond to membrane invaginations (arrows, **Fig. 1C**). Transmission EM on ultrathin 2D sections further demonstrated the presence at the midbody (MB) of small (60-100 nm) plasma membrane invaginations with typical morphology of caveolae (arrow, **Fig. 1D**).

**Figure 1.**
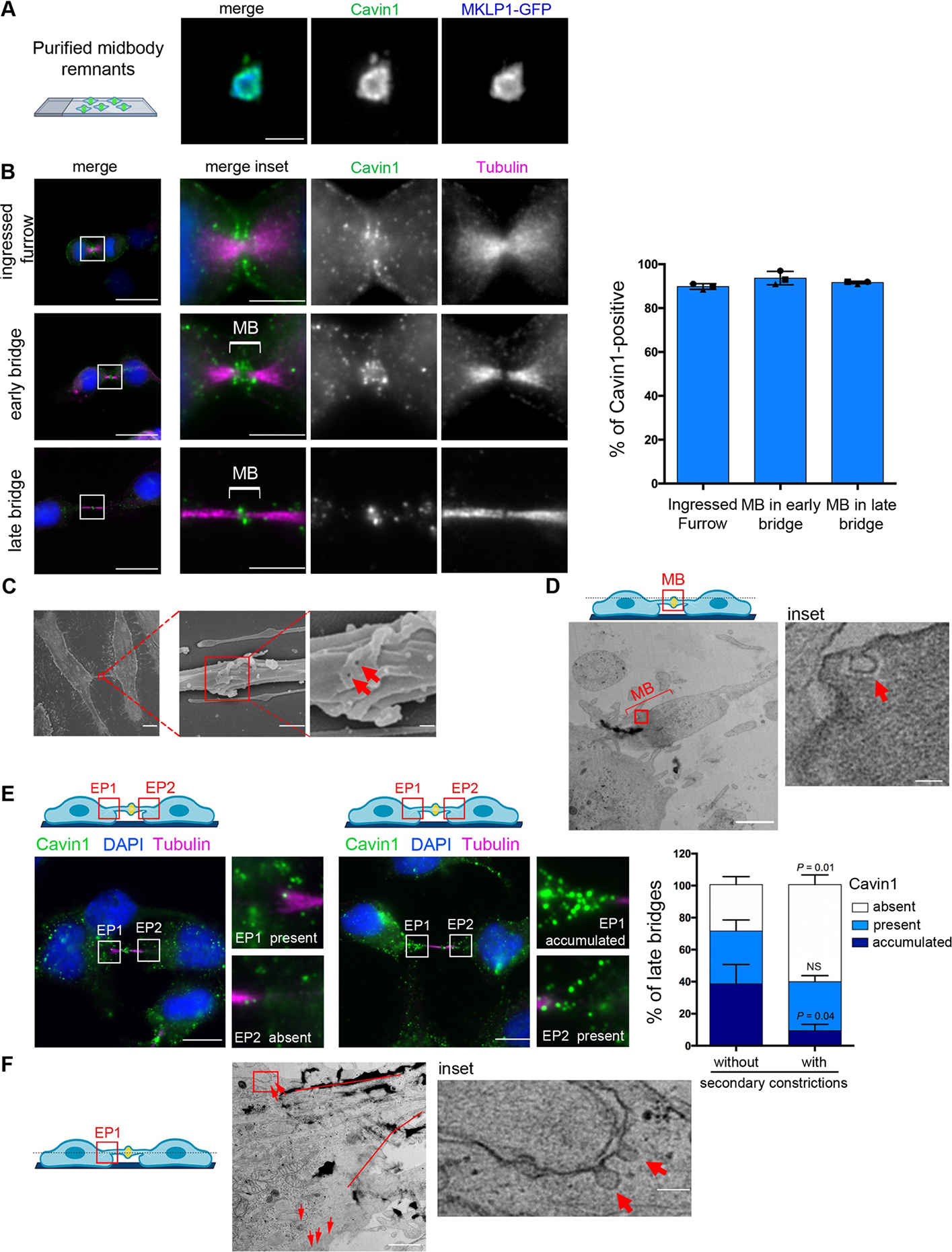
Caveolae are present at the midbody and at the Entry Points (EPs) during cytokinesis. **(A)** Purified midbody remnants from HeLa cells that stably express MKLP1-GFP were immunostained for endogenous Cavin1. Scale bar, 2 µm. **(B)***Left panels:* Localization of endogenous Cavin1 in HeLa cells stained with *α*-tubulin + acetylated- tubulin (hereafter “Tubulin”) and DAPI at the indicated stage of cytokinesis. Scale bars, 10 µm (general views), 2 µm (insets). *Right panel:* Percentage of the indicated structures positive for endogenous Cavin1. Mean ± SD, n> 20 cells per condition. N = 3 independent experiments. MB: midbody. **(C)** Scanning electron microscopy (SEM) pictures of an intercellular bridge (left) and midbody at different magnifications of the indicated boxed regions (successive insets). Red arrows point to membrane invaginations at the midbody. Scale bars, 10 µm (left), 500 nm (middle) and 100 nm (right). **(D)***Upper panel:* Scheme highlighting the position of the midbody (MB) within the intercellular bridge (approximately 5 µm above the substrate). *Bottom panels:* Transmission electron microscopy (TEM) pictures of a midbody (MB, electron dense part within the ICB) with the inset from the indicated boxed region showing caveolae (red arrow). Scale bars, 10 µm (left) and 100 nm (inset). **(E)***Upper panels:* Schemes highlighting the localization of the entry points (EP1, EP2), which correspond to the regions (red squares) at the intercellular bridge/cell interfaces. *Lower panels:* Two examples of endogenous staining for Cavin1, together with Tubulin and DAPI, in late bridges. Insets show different categories of Cavin1 localization at EPs: accumulated, present or absent, as indicated. Scale bars, 10 µm. *Right panel:* Percentage of late bridges with/without secondary constriction where Cavin1 is accumulated, present or absent at EPs, as indicated. Mean ± SD, n> 30 cells per condition. N = 3 independent experiments. One-sided Student’s *t* tests. NS nonsignificant (P > 0.05). **(F)***Left panel:* Scheme highlighting the localization of an EP region chosen for EM analysis. *Right panels:* TEM pictures of an EP region with the membrane of the ICB highlighted with red lines and caveolae marked with red arrows. The inset shows caveolae (red arrows) of the indicated boxed region. Scale bars, 10 µm and 100 nm (inset). In (D, F) the electron-dense (black) spots represent deposits of contrasting agents.

After furrow ingression, we also frequently observed an accumulation of dotty Cavin1- and Cav1-positive structures at one or both sides of the ICB (**Fig. 1E and S1E**). These flanking regions of the ICB correspond to the interface between the ICB and the daughter cell, and we thus named them “entry points” (EPs). Transmission EM confirmed that caveolae were present at the plasma membrane at the EPs (**Fig. 1F**). In late cytokinetic bridges, secondary constrictions with decreased microtubule staining are found ∼1 μm away from midbody and correspond to the tip of the ESCRT-III cone that constrict at the presumptive abscission site (*4, 39, 43*) (see **Fig. S1A**). Interestingly, the accumulation of caveolae at EPs was observed in approximately 40% of late ICBs without secondary constrictions but only in 10% of ICBs with secondary constrictions (which correspond to ICBs temporally closer to abscission) (**Fig. 1E and S1E**). This indicates that the number of caveolae at the EPs decreases as cells progress towards abscission. Altogether, we conclude that *bona fide* caveolae are present at the midbody and transiently at the ICB/cell interface after furrow ingression.

### Cavin1 and Cav1 dynamically colocalize at the midbody and at the ICB/cell interface

To reveal the dynamics of the caveolae during cytokinesis, we monitored their localization in live cells using time-lapse spinning-disk confocal microscopy in cells expressing both Cav1-RFP and Cavin1-NG (mNeonGreen). As shown in **Fig. 2A** and **Movie 1**, punctate structures positive for both Cavin1 and Cav1 localized at the ingressed furrow (190 min before abscission, time -190 min). As the ICB matured, pools of Cavin1 and Cav1 were found at the midbody (**Fig. 2A**, bracket, times -120, -70 and 0 min) and accumulated at EPs (**Fig. 2A**, squared regions, time -160 min). Interestingly, quantification of Cavin1-NG intensity at EPs (**Fig. 2B, Fig. S2A** and **Movie 2**) revealed that 1) one EP is consistently more labeled than the other, suggesting an asymmetry in caveolae localization at EPs; 2) Cavin1 intensity starts to decrease at both EPs approximately 100 minutes before abscission, with a sharper decrease 50 min before abscission; 3) abscission occurs approximately 20 minutes after both EPs have reached their minimal Cavin1 intensity. Since ESCRT-III starts to polymerize as a cone from the midbody toward the abscission approximately 30 min before abscission (*43*) (see **Fig. S1A**), ESCRT-III polymerization is thus concomitant with low levels of Cavin1 at EPs. Importantly, Cavin1-positive dots detected in live cell imaging were almost always positive for Cav1 (**Fig. 2A**, lower panel) indicating that they correspond to caveolae (*20℃22*), which is consistent with our EM data in fixed cells.

**Figure 2.**
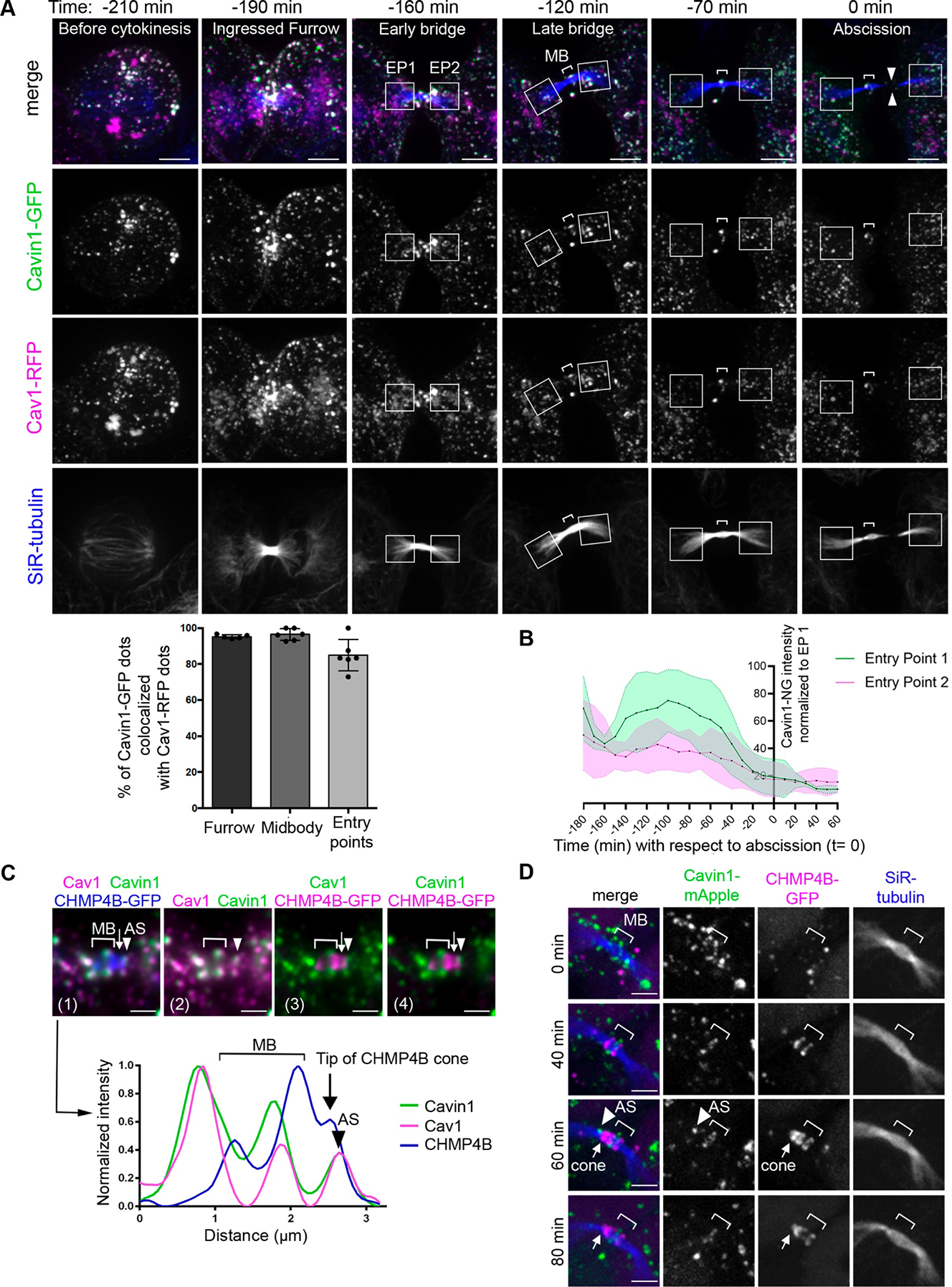
Cavin1 and Cav1 dynamically colocalize at the midbody, the EPs and the abscission site. **(A)** *Upper panels:* Snapshots of a spinning-disk confocal microscopy movie of cells co- expressing Cavin1-GFP and Cav1-RFP. SiR-Tubulin was added for microtubule visualization. Squares delimit each EP and brackets indicate the position of the midbody (MB). Facing arrowheads mark the abscission site. Time 0 indicates the time of abscission. Note the progressive disappearance of Cavin1/Cav1 in EP regions as cells progress to abscission. Scale bars, 5 µm. *Lower panel:* Percentage of Cavin1-GFP dots colocalizing with Cav1-RFP dots at the furrow, midbody and EPs. Mean ± SD, n= 6 movies of dividing cells. **(B)** Intensity of Cavin1-GFP staining as a function of time at EP1 and EP2 in n= 10 cells throughout cytokinesis (see snapshots of a representative video in Fig. S2A). The brightest EP was defined as EP1 and the values of EP1 (green) and EP2 (magenta) were normalized to the maximum intensity observed in EP1 during cytokinesis for each cell analyzed. For comparison, all movies were registered with respect to the time of abscission (t= 0). Mean ± SD, n= 10 movies. **(C)** *Upper panels:* Endogenous staining for Cav1 and Cavin1 in cells stably expressing CHMP4B-GFP, with a zoom centered on the ICB (1). Same bridge with only Cav1 and Cavin1 (2), Cav1 and CHMP4B (3) and Cavin1 and CHMP4B (4) channels displayed. Bracket: midbody (MB). Arrow: CHMP4B-GFP polymerizing cone. Arrowhead: abscission site where Cavin1/Cav1 dots co-localize at the tip of the CHMP4B cone. *Lower panels:* Intensity profiles along the bridge of the merged picture (1) with matched colors and symbols. Scale bars, 1 µm. **(D)** Snapshots of a spinning-disk confocal microscopy movie of a cell expressing Cavin1- mApple and CHMP4B-GFP. SiR-Tubulin was added for microtubule visualization. Bracket: midbody (MB). Arrow: CHMP4B-GFP polymerizing cone. Arrowhead: abscission site (AS). Note that the green Cavin1 dot at the tip of the CHMP4B cone disappears while the cone constricts the microtubules at the future abscission site. Scale bars, 2 µm.

Beside the midbody and EPs, we also noticed the presence of discrete Cavin1- and Cav1-positive punctate structures at the future abscission site (**Fig. S2B**, arrowheads). These structures contained both Cavin1 and Cav1, and were localized at the distal tip (**Fig. 2C**, arrowhead) of the ESCRT-III cone labelled by CHMP4B (**Fig. 2C**, arrow). Remarkably, time lapse microscopy demonstrated that this pool of caveolae disappeared prior to abscission, while the ESCRT-III cone polymerized toward the abscission site (100 % of cases, n= 13 movies and **Fig. 2D**).

In summary, caveolae are found at the midbody throughout cytokinesis. They are also found at the ICB/cell interface (*i.e.* entry points) and at the tip of the ESCRT-III machinery, and progressively disappear from these locations as cells progress toward abscission.

### Loss of caveolae impairs abscission and ESCRT-III localization

To investigate the function of caveolae in cytokinesis, we took advantage of a HeLa cell line KO for Cavin1 that we recently generated using CRISPR-Cas9 strategy (*44*). Westernblots confirmed the absence of Cavin1 protein expression in these cells (**Fig. 3A**). In addition, we observed the loss of Cavin1 dots by immunofluorescence (**Fig. S3A**). As expected, this was accompanied with a strong reduction of Cav1-positive dots upon Cavin1 KO (**Fig. S3B**), since Cavin1 and Cav1 protein levels are correlated (*45, 46*). Using transmission EM, we did not detect the presence of caveolae in Cavin1 KO cells, contrary to control KO cells (**Fig. 3B**, arrows). We conclude that caveolae are absent in the Cavin1 KO cells that we generated.

**Figure 3.**
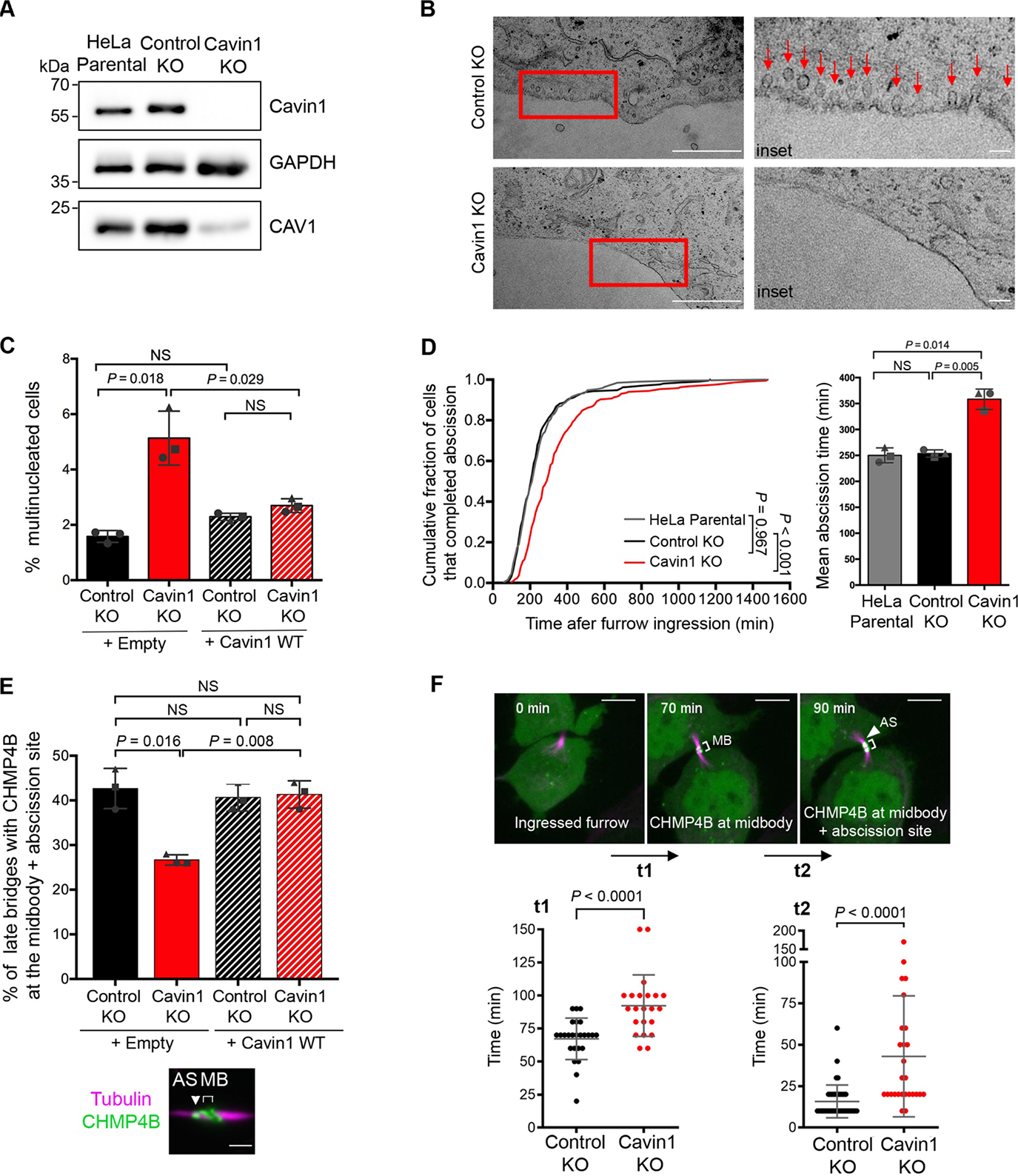
Depletion of Cavin1 impairs ESCRT-III localization and abscission. **(A)** Lysates of parental HeLa, Control KO and Cavin1 KO cells were blotted for Cavin1, Cav1 and GAPDH (loading control). **(B)** TEM micrographs of ultrathin sections of Control KO and Cavin1 KO cells. Insets from the boxed regions highlight the presence of caveolae (red arrows) in Control KO cells and absence of caveolae in Cavin1 KO cells (at least 10 individual cells were analyzed). Scale bars, 1 µm and 100 nm (insets). **(C)** Percentage of Bi-/Multi-nucleated cells in Control KO and Cavin1 KO cells transfected with the indicated plasmids. Mean ± SD, n> 500 cells per condition. N = 3 independent experiments. **(D)** *Left panel:* Cumulative distribution of the fraction of cells that completed abscission, measured by phase-contrast time lapse microscopy in HeLa Parental, Control KO and Cavin1 KO cells. *P* values from nonparametric and distribution-free Kolmogorov– Smirnov [KS] tests. *Right panels:* Mean abscission times (min) ± SD in indicated cells, with n> 60 cells per condition. N = 3 independent experiments. **(E)** The proportion of late cytokinetic bridges with endogenous CHMP4B localized both at the midbody and at the abscission site (see representative image) was quantified in Control KO and Cavin1 KO cells transfected with indicated plasmid. Mean ± SD, n > 30 cells per condition. N = 3 independent experiments. Arrowhead: abscission site (AS); midbody (MB): bracket. Scale bar, 2 µm. **(F)** *Upper panels:* Control KO and Cavin1 KO cells stably expressing CHMP4B-GFP were recorded by spinning-disk confocal time-lapse microscopy. The time taken from cleavage furrow ingression (t= 0 min) to CHMP4B-GFP recruitment at the midbody defines t1; and time from CHMP4B-GFP at the midbody to CHMP4B-GFP at midbody + abscission site defines t2. Brackets: midbody (MB). Arrowhead: CHMP4B polymerizing toward the future abscission site. Scale bars, 5 µm. *Lower panels:* Times (min) ± SD for t1 and t2 in Control KO and Cavin1 KO. n> 23 cells per condition. N = 4 independent experiments. In (C), (D) and (E): Two-sided Student’s *t* tests. In (F): Unpaired Student’s *t* tests. NS non- significant (P > 0.05).

Most noteworthy, the lack of caveolae was associated with a 3-fold increase in the number of binucleated cells in the Cavin1 KO cell population (**Fig. 3C**), indicating cytokinetic defects. Furthermore, using time-lapse phase contrast microscopy, we observed that abscission was delayed in Cavin1 KO cells compared to control cells (**Fig. 3D**). Importantly, re-expression of Cavin1 in the KO cells fully rescued the cytokinetic defects, ruling out off target artifacts (**Fig. 3C and S3C**). Similarly, depletion of Cav1 by siRNAs resulted in the loss of Cav1-positive structures in the ICB (**Fig. S3D**) and delayed abscission as well (**Fig. S3E**).

To understand the origin of the cytokinetic defects induced by the loss of caveolae, we next investigated the ESCRT scission machinery by monitoring the key ESCRT-III subunit CHMP4B. In fixed cells, Cavin1 KO led to a decrease in the proportion of ICBs with CHMP4B both at the midbody and at the abscission site (**Fig. 3E**). This suggested a defect in ESCRT-III assembly at the abscission site upon removal of caveolae. As expected, the abnormal localization of CHMP4B was fully rescued by re-expressing Cavin1 (**Fig. 3E**). Spinning-disk confocal microscopy further demonstrated delayed arrival of CHMP4B-GFP both at the midbody (t1, **Fig. 3F**) and at the abscission site (t2, **Fig. 3F**) upon Cavin1 KO. These results indicate that caveolae are required for the proper assembly of ESCRT-III at the midbody and at the abscission site. Altogether, this explains, at the mechanistic level, the abscission defects observed upon caveolae loss.

### Caveolae limit acto-myosin II contractility at the ICB/cell interface and control ICB tension during cytokinesis

High tension in ICBs during cytokinesis delays ESCRT-III polymerization (*17*). Since membrane tension was found to contribute to half of the bridge tension (*17*) and caveolae can buffer membrane tension in interphase cells (*25, 47*), we hypothesized that the loss of caveolae may increase ICB tension during cytokinesis. Changes in ICB tension can be measured by laser ablation. Indeed, when the ICB is transversally cut by a UV- laser (**Fig. 4A**), the initial speed of retraction was previously shown to be proportional to the forces exerted by the daughter cells on their ICB (*17*). Remarkably, using this approach, we found a two-fold increase of ICB tension in Cavin1 KO cells compared to control KO cells (**Fig. 4B and Movie 3**). These results demonstrate that caveolae are functionally required to limit the ICB tension during cytokinesis.

**Figure 4.**
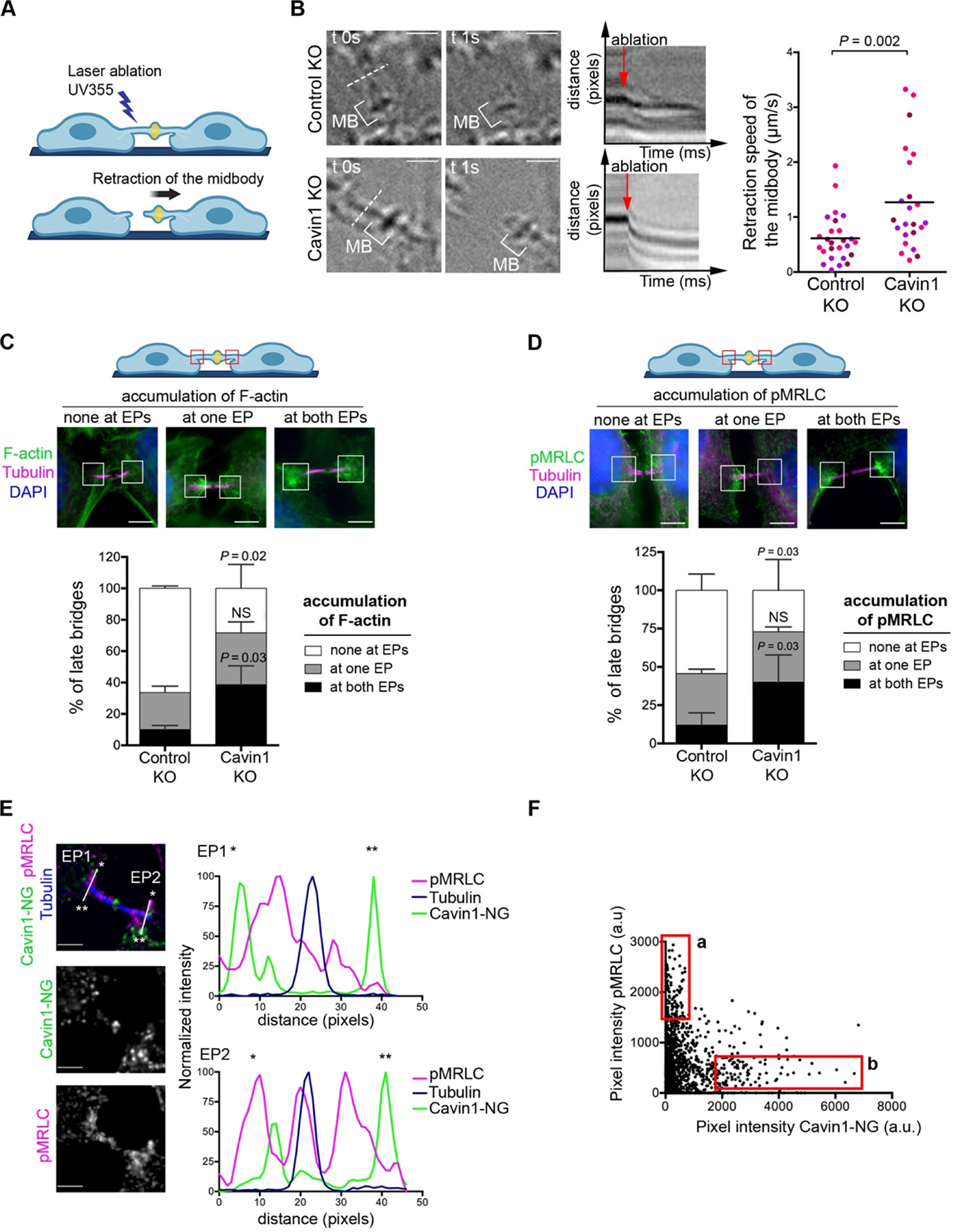
Caveolae limit acto-myosin II at the Entry Points and control ICB tension during cytokinesis. **(A)** Scheme illustrating the laser ablation experiment. A laser-induced ablation of a tensed ICB results in a retraction of the midbody toward the side opposite to the cut. **(B)** *Left Panels:* Snapshots of ICBs from Control KO and Cavin1 KO cells, before (t0) and after the laser ablation (t0 + 1 sec). The corresponding kymographs are displayed, with red arrows marking the moment of ablation. Dotted line: position of the laser cut. Brackets: midbody (MB). Scale bars, 2 µm. *Right Panel:* Retraction speed (µm/s) of midbodies in Control KO and Cavin1 KO ablated bridges. Horizontal bar: Mean, n= 23-24 ICBs from N = 3 independent experiments (represented with different colors). Unpaired Student’s *t* tests. **(C)** *Upper Panels:* The accumulation of endogenous F-actin staining (fluorescent phalloidin) in EPs in late cytokinetic bridges was classified in three categories: none at EPs, at one EP and at both EPs, as shown in representative images. Scale bar, 5 µm. *Lower Panels:* Percentage of late bridges for each F-actin category in Control KO and Cavin1 KO cells. Mean ± SD, n > 20 cells per condition. N = 3 independent experiments. Paired Student’s *t* tests. NS nonsignificant (P > 0.05). **(D)** *Upper Panels:* The accumulation of endogenous pMRLC staining in EPs of late cytokinetic bridges was classified in three categories: none at EPs, at one EP and at both EPs, as shown in representative images. Scale bars, 5 µm. *Lower Panels:* Percentage of late bridges for each pMRLC category in Control KO and Cavin1 KO cells. Mean ± SD, n > 30 cells per condition. N = 3 independent experiments. Paired Student’s *t* tests. NS nonsignificant (P > 0.05). **(E)** *Left panels:* Endogenous pMRLC and Tubulin staining in a Cavin1-NeonGreen stable cell line, with a zoom centered on the ICB. Scale bars, 2 µm. *Right panels:* intensity profiles of the EP1 and EP2 regions, as in indicated in the merged picture with matched colors symbols. **(F)** Pixel intensity of pMRLC (y axis) and Cavin1-NG (x axis) at the whole EP1 and EP2 from (E). Box a: high pMRLC/low Cavin1 intensity pixels; Box b: high Cavin1/low pMRLC intensity pixels.

Tension measured in the ICB can result from an entangled combination of membrane tension and cell contractility transmitted to the ICB by the acto-myosin system (*48*). We thus examined the activity of myosin II by measuring the levels of phosphorylated Myosin II Regulatory Light Chain (pMRLC) and F-actin localization upon Cavin1 depletion. First, we observed that the proportion of cytokinetic cells with F-actin accumulation at both EPs increased in Cavin1 KO cells (**Fig. 4C**). Conversely, the proportion of cytokinetic cells without F-actin accumulation at either EPs decreased in Cavin1 KO cells (**Fig. 4C**). Second, as seen for F-actin, the proportion of cytokinetic cells with accumulation of endogenous pMRLC at both EPs increased in Cavin1 KO cells (**Fig. 4D**). Conversely, the proportion of cytokinetic cells without pMRLC accumulation at either EPs decreased in Cavin1 KO cells (**Fig. 4D**). However, neither the total levels of MRLC nor the percentage of activated myosin II (ratio pMRLC/total MRLC) was changed in Cavin1 KO cells synchronized in cytokinesis (**Fig. S4A**). Altogether, the loss of caveolae results in a higher proportion of ICBs displaying F-actin and activated myosin II in the EP regions.

Interestingly, in wild type cells, we observed that structures with high levels of activated myosin II and high levels of Cavin1 were inversely correlated (and often mutually exclusive) within EP regions: high intensity Cavin1-positive structures (**Fig. 4E** green peaks and **Fig. 4F box b**) were negative or weak for pMRLC (**Fig. 4E** magenta peaks and **Fig. 4F box b**) and vice-versa (**Fig. 4E and Fig. 4F box a**). This suggests that the presence of caveolae locally restricts myosin II activation at EPs during cytokinesis, a hypothesis consistent with increased pMRLC at EPs found in the absence of Cavin1.

Altogether, we conclude that the presence of caveolae both limits the presence of F-actin/active myosin II at the cell/ICB interface and the ICB tension during cytokinesis.

### Increased ICB tension is responsible for the cytokinetic defects induced by the loss of caveolae

Our data raised the possibility that, upon Cavin1 depletion, the local increase in F-actin and activated myosin II at EPs could contribute to the observed increase in ICB tension, thus explaining the defects in ESCRT-III assembly and the abscission delay. To test this hypothesis, we treated Cavin1 KO cells with Y27632, a pharmacological inhibitor of ROCK, a kinase that activates myosin II through direct phosphorylation of MRLC. Upon ROCK inhibition, the increase in the proportion of ICBs with both EPs positive for pMRLC observed in Cavin1 KO cells was abolished (**Fig. 5A**). Concomitantly, ROCK inhibition restored ICB tension to normal levels in Cavin1 KO cells (**Fig. 5B**), suggesting that activated myosin II at EPs contributes to the ICB tension increase measured upon Cavin1 depletion. Remarkably, the treatment of Cavin1 KO cells with Y27632 was also sufficient to restore the correct assembly of ESCRT-III at the abscission site (**Fig. 5C**). Furthermore, reducing membrane tension by incubating the cells in a moderately hyper-osmotic medium (400 mOsm) also restored the proper assembly of ESCRT-III at the abscission site in Cavin1 KO cells (**Fig. S4B**). These results show that reducing either membrane tension or acto-myosin II-dependent contractility is sufficient to correct the abnormal localization of ESCRT-III at the abscission site observed upon the loss of caveolae. Accordingly, Y27632 treatment also rescued the abscission defects observed in Cavin1 KO cells (**Fig. 5D**). Noteworthy, in wild type cells, we observed that the polymerization of ESCRT-III at the abscission site occurs concomitantly with a decrease in the accumulation of pMRLC at EPs (**Fig. S4C**). This suggests that during normal cytokinesis, the drop in ICB tension known to promote abscission (*17*) depends on the decrease of acto-myosin II contractility at EPs.

**Figure 5.**
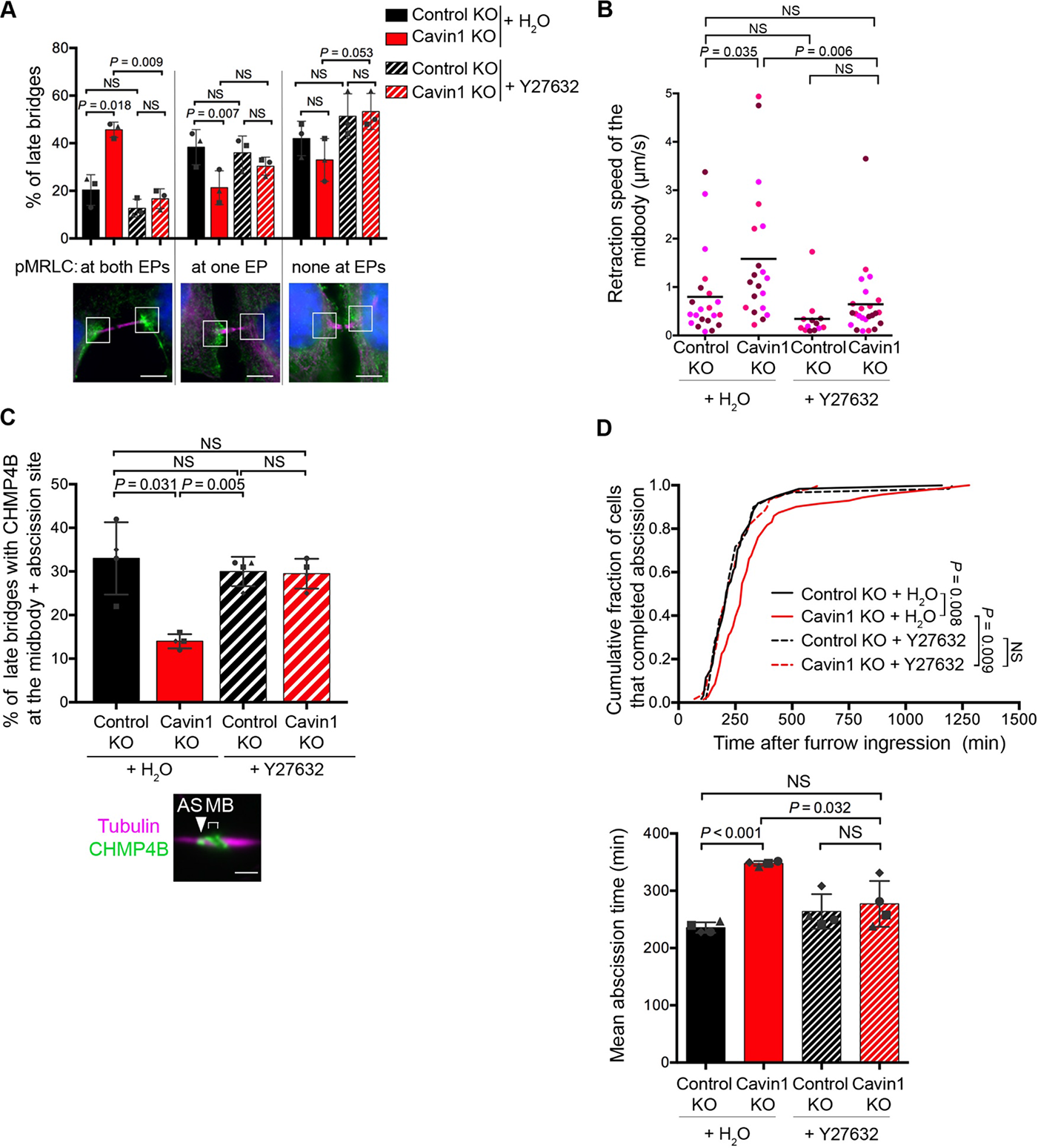
Increased ICB tension upon Cavin1 depletion is responsible for the cytokinetic defects. **(A)** Quantification of pMRLC localization at the EPs as in Fig. 4D (same representative pictures, Scale bars, 5 µm) in Control KO and Cavin1 KO cells, treated or not with the ROCK inhibitor Y27632. Horizontal bar: Mean ± SD, n> 30 cells per condition. N = 3 independent experiments (represented with different colors). **(B)** Retraction speed (µm/s) of midbodies in Control KO and Cavin1 KO ablated bridges, treated or not with the ROCK inhibitor Y27632. Horizontal bar: Mean, n> 12 ICBs from N = 3 independent experiments. Unpaired Student’s *t* tests. NS non-significant (P > 0.05). **(C)** CHMP4B localization was quantified as in Fig. 3E (same representative picture, Scale bar, 2 µm) in Control KO and Cavin1 KO cells, treated or not with the ROCK inhibitor Y27632. Mean ± SD, n> 30 cells per condition. N = 3 independent experiments. **(D)** *Upper panel:* Cumulative distribution of the fraction of cells that completed abscission, measured by phase-contrast time lapse microscopy in Control KO and Cavin1 KO cells, treated or not with the ROCK inhibitor Y27632. *P* values from nonparametric and distribution-free Kolmogorov–Smirnov [KS] tests. NS non-significant (*P* > 0.05). *Lower panel:* Mean abscission times (min) ± SD in indicated cells, with n> 60 cells per condition. N = 4 independent experiments. In (A), (C) and (D): paired Student’s *t* tests. NS non-significant (*P* > 0.05).

We conclude that the increase of tension in the ICB induced by the absence of caveolae is the cause of the defects in ESCRT-III assembly, which in turn impairs normal abscission. Therefore, caveolae are functionally required to promote ESCRT-III polymerization and successful abscission by limiting intercellular bridge tension during cytokinesis.

## DISCUSSION

The decrease in ICB tension after furrow ingression is one of the most upstream mechanical determinants that controls the assembly of the abscission machinery. Yet, no regulator of ICB tension has been reported so far. Here, we establish the first functional role of caveolae in cell division, which were found to buffer ICB tension to promote cytokinetic abscission.

Our recent proteomic analysis of MBRs purified by flow cytometry revealed the presence or the enrichment of several structural and regulatory components of caveolae (Cav1, Cavin1, Cavin3, EHD2, PACSIN2-3). The compilation of other proteomic studies (*42, 49, 50*) of either intercellular bridges or MBRs released into the culture medium, confirmed this finding. This led us to hypothesize that caveolae could play an active role during cytokinesis. Indeed, proteins involved either in furrow ingression or in late cytokinetic stages often accumulate at the midbody and can be identified by mass spectrometry in purified MBRs (*43*). This recently allowed us to reveal the role of ALIX/syntenin/Syndecan-4 and BST2/tetherin in abscission and post-abscission MBR capture, respectively (*43, 51*).

While a role of caveolae in cytokinesis has not been reported so far, the structural caveolae component Cav1 had been previously localized at the furrow (*36, 37*). We confirmed and extended these observations using endogenous staining for both Cav1 and Cavin1 (Fig. 1B, S1D) as well as fluorescently-tagged proteins (Fig. 2A). We also revealed the presence of caveolae components 1) at the midbody itself, 2) at the “entry points” (EPs), which correspond to the interfaces between the ICB and the daughter cells (Fig. 1E, S1E, 2A, S2A) and 3) at the tip of the ESCRT-III cone while polymerizing toward the abscission site (Fig. 2C-D). The extensive colocalization between Cavin1 with Cav1 in these structures (Fig. 2A), together with EM data (Fig. 1D,F) demonstrated that these are *bona fide* caveolae present at the plasma membrane.

What could determine the localization of caveolae at these specific locations during cytokinesis? Intense recycling of caveolae components during mitotic exit and telophase has been reported (*35*), and it is thus possible that this contributes to the localization of caveolae at the cytokinetic furrow, EPs and midbody. Alternatively, individual caveolae could diffuse from the cell body to the furrow, EPs and midbody within the plane of the plasma membrane. Finally, caveolae could as well form *in situ* due to the peculiar lipid composition of the furrow and midbody membrane that is known to promote caveolae formation in non-dividing cells (*52–57*). We also noticed that caveolae are particularly abundant at EPs when the plasma membrane adopts a funnel shape (as in Fig. 1E, “EP1 accumulated” right panels), possibly indicating that specific curvature might favor caveolae formation or retention. Regarding the intriguing presence of caveolae at the tip of the ESCRT-III cone, we noticed that the ESCRT-III subunit CHMP4B had been found in complex with several caveolae components (Cavin1, Cav1 and PACSIN3) using mass spectrometry (*58*). This could provide a molecular connection explaining the remarkable coupling between caveolae and ESCRT-III at the abscission site.

What is the role of caveolae during cytokinesis? Following furrow ingression, daughter cells exert nano Newton-range pulling forces on the ICB that must be reduced to favor abscission (*17*). Accordingly, high tension inhibits ESCRT-III polymerization at the abscission site during cytokinesis (*17*) and prevents membrane scission in reconstitution assays (*59*). Our results point to a key role of caveolae in limiting ICB tension, which in turn promotes ESCRT-III polymerization and consequently abscission. Indeed, the depletion of Cavin1 results in 1) a 2-fold increase in ICB tension (Fig. 4B); 2) a delay in ESCRT-polymerization at the midbody and at the abscission site (Fig. 3E-F) and 3) a delay in abscission (Fig. 3D) or cytokinetic failure (Fig. 3C). In addition, Cav1 depletion leads to similar defects in cytokinesis (Fig. S3E). Importantly, the increase in ICB tension upon caveolae loss is the cause of the cytokinetic failure, since decreasing membrane tension through a moderate hyperosmotic shock (Fig. S4B) or by decreasing acto-myosin II-dependent contractility through ROCK inhibition (Fig. 5C) restores the proper polymerization of ESCRT-III at the abscission site in Cavin1 KO cells. Remarkably, ROCK inhibition also rescues the abscission defects in these cells (Fig. 5D). To our knowledge, caveolae thus appear as the first regulator of ICB tension during cytokinesis.

How could caveolae control ICB tension during cytokinesis? ICB tension results from a combination of membrane tension and cortical tension due to acto-myosin II- dependent contractility exerted on the ICB (*17, 48*). We envision two, non-mutually exclusive, mechanisms to explain how caveolae might control both components of ICB tension.

First, the pool of caveolae present at the EPs could contribute to the decrease in membrane tension at the ICB by flattening out (**Fig. 6**, wild type cells), as shown previously in non-dividing cells (*24, 48*). Consistently, using both fixed cells (Fig. 1E, S1E) and 10-min frequency time-lapse microscopy (Fig. 2A, 2B and S2A), we found that caveolae progressively disappeared from EPs as cells progressed through abscission. The accumulation of caveolae precisely at the interface between the ICB and the cell body would constitute a membrane reservoir ideally located for reducing membrane tension at the ICB. The caveolae-mediated membrane tension reduction would then promote ESCRT-III polymerization at the abscission site and thus abscission. In agreement with the idea that caveolae flattening at EPs reduces ICB tension, it is known that ESCRT-III starts to polymerize as a cone at the presumptive abscission site approximately 30 min before abscission occurs, and this corresponds to the time when caveolae at the EPs intensity reaches a minimum (Fig. 2B). Interestingly, previous quantitative measurements showed that the tension at the ICB also diminishes at that time (*17*). Beside caveolae flattening at EPs, caveolae flattening at the plasma membrane of the daughter cells, outside the EP regions, could also contribute to the reduction of ICB tension that precedes abscission. In addition, the flattening of the caveolae localized at (or close to) the tip of the ESCRT-III cone (Fig. 2C-D) could also help the constriction of the plasma membrane at the abscission site by locally reducing membrane tension.

**Figure 6.**
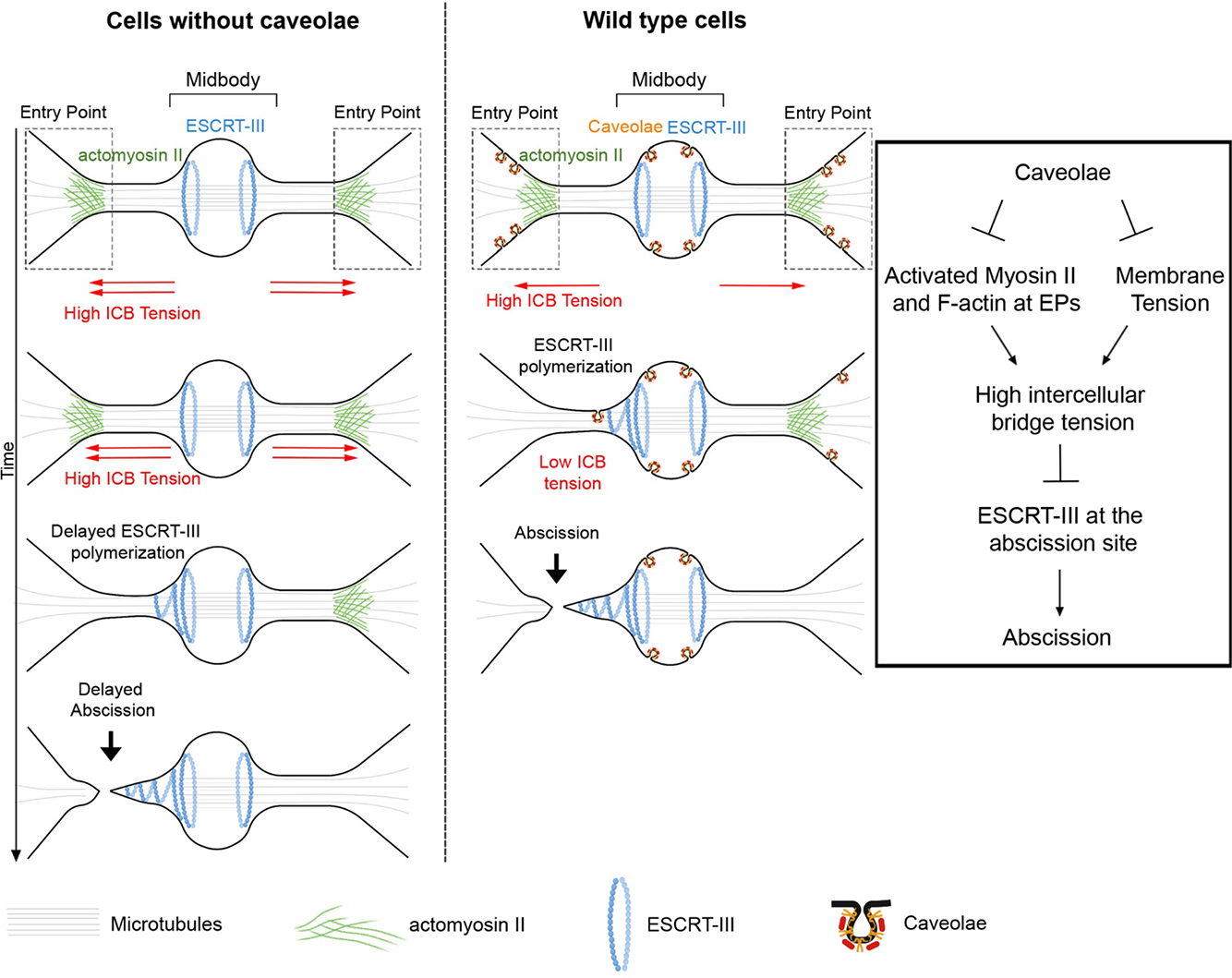
Model: Caveolae buffer ICB tension by limiting the F-actin/myosin II- dependent contractility and membrane tension, thereby promoting ESCRT-III polymerization and successful abscission. *Left panel:* situation in cells without caveolae (e.g. Cavin1 KO cells). Right panels: situation in cells with caveolae (wild type cells). *Box:* proposed mechanism for the control of abscission by caveolae.

Second, caveolae at EPs could also negatively regulate acto-myosin II-dependent contractility at or close to the ICB. Indeed, depleting Cavin1 leads to an increase of the proportion of ICBs with activated myosin II at EPs, associated with a 2-fold increase in ICB tension (Fig. 4B, D). This regulation appears local, since global myosin II activation during cytokinesis is unchanged after caveolae depletion (Fig. S4A). Furthermore, pharmacological inhibition of ROCK in Cavin1 KO cells enabled to restore both normal pMRLC patterns at the EPs as well as normal ICB tension (Fig. 5A-B). This suggests that the abnormally high ICB tension observed after caveolae loss results from the local increase in acto-myosin II activity at the EPs. How the presence of caveolae at EPs seen in wild type cells locally regulates this previously undescribed pool of activated myosin II to restrict ICB tension is not known and would require future investigation. Nonetheless, the inverse correlation between high levels of activated myosin II and high levels of Cavin1 at EPs is remarkable (Fig. 4E-F) and might involve complex biochemical reciprocal inhibitory signals between caveolae and the acto-myosin system. Interestingly, there is increasing evidence that caveolae regulate both membrane tension and acto-myosin II activity at the plasma membrane (*32, 60–63*), in particular during rear-to-front retraction during durotaxis-mediated migration (*62*) and in cell-cell contacts involved in melanin pigment transfer from melanocytes to keratinocytes (*63*).

Furthermore, caveolae have been recently implicated in the negative regulation of F-actin polymerization and thereby intercellular tension through adherent junctions between non-dividing epithelial cells (*32*). The parallel with the role of caveolae as a negative regulator of ICB tension during cell division is striking, although the functional consequence —ESCRT-III polymerization and membrane scission— is very different. Thus, the local regulation of acto-myosin-dependent contractility coupled to the buffering of membrane tension might represent a general function of caveolae.

In summary, we propose that the presence of caveolae at EPs and at the abscission site buffers membrane tension and limits myosin II activation at the ICB in wild type dividing cells (Fig. 6, right panels). This promotes the decrease of the ICB tension, thus favoring the polymerization of ESCRT-III at the abscission site to allow abscission. In cells without caveolae, the lack of the caveolae membrane reservoir and the increased activation of myosin II at EPs leads to an increase of ICB tension that is detrimental for ESCRT-III polymerization and abscission (Fig. 6, left panels). This could be particularly relevant in tumorigenesis, since caveolae both promote elimination of cancer cells through extrusion (*32*) and prevents the formation of binucleated cells (this study), a well-established starting point for tumor formation (*64, 65*). The unexpected connection between caveolae and the ESCRT machinery should also stimulate investigation on how caveolae might control key other ESCRT-III-dependent processes beyond cell division, such as the budding of enveloped viruses and plasma membrane repair.

## MATERIALS AND METHODS

### Cell Culture

HeLa CCL-2 (ATCC), HeLa Control KO (HeLa CCL-2 transfected with a plasmid encoding Cas9 without gRNA described in (*44*)), HeLa Cavin1 KO (HeLa CCL-2 transfected with a plasmid encoding Cas9 and a gRNA targeting *Cavin1* described in (*44*)) and HeLa CHMP4B-GFP described in (*43*) were grown in Dulbecco’s Modified Eagle Medium (DMEM) GlutaMax (#31966 Gibco, Invitrogen Life Technologies). All culture media were supplemented with 10 % fetal bovine serum (FBS, #500105R1 Dutscher) and 1 % Penicillin/Streptomycin (#15140122 Gibco) and cultured in 5 % CO_2_ condition at 37 °C. CHMP4B-GFP stable cell lines were generated by lentiviral transduction of HeLa Control KO and Cavin1 KO cells and selected by flow cytometry (FACS) sorting. For the ROCK inhibitor experiments, HeLa Control KO and Cavin1 KO were treated with Y27632 (Calbiochem) at 20 μM overnight (Fig. 5B, 5D) or 50 μM for 3h (Fig. 5A, 5C).

### Plasmid constructs

Cavin1-GFP and Cav1-RFP were described previously in (*66*). To construct Cavin1- mApple, the Cavin1 CDS was subcloned into Gateway pENTR plasmids. The transient expression vector was generated by LR recombination (Thermo Fisher) of Cavin1-pENTR and pDEST mApple N1.

### Cell transfections and siRNAs

For live cell imaging, HeLa cells, were transfected with 200 ng of plasmids for 48 h using X-tremeGENE 9 DNA reagent (Roche, Fig. 2A), or electroporated with 2 μg of plasmid using Lonza’s nucleofection protocol (Fig. S2A). HeLa CHMP4B-GFP cells were transfected with 80 ng Cavin1-mApple for 48h using X-tremeGENE 9 DNA reagent (Fig. 2D). For the rescue experiments, Control KO and Cavin1 KO cells were transfected with 200 ng of plasmids (Fig. 3C, S3C) or 120 ng of plasmid for 24h (Fig. 3E) for 24h using X- tremeGENE 9 DNA reagent (Roche).

Depletion of Cav1 was achieved by RNAi using siRNAs with the following sequence: 5’ GCAUCAACUUGCAGAAAGA 3’ (Merck), and Control siRNA Luciferase: 5’ CGUACGCGGAAUACUUCGA 3’ (Merck). HeLa cells were transfected on two consecutive days with 20 nM Cav1 siRNAs using Lipofectamine RNAiMAX (Invitrogen), following the manufacturer’s instructions and recorded 3 days later.

### Western Blots

siRNAs-treated HeLa cells and Control KO and Cavin1 KO cells were directly lysed in 1x Laemmli (Bio-Rad Laboratories) with Benzonase nuclease (#70664 Millipore) and migrated in 10 % SDS-PAGE gels (Bio-Rad Laboratories). Control KO and Cavin1 KO cells used in the rescue experiments were detached with trypsin (Gibco), collected and lysed with NP-40 extract buffer (50 mM Tris, pH 8, 150 mM NaCl, 1 % NP-40) containing protease inhibitors (#11873580001 Roche) and 30 µg of lysate was migrated in 10 % SDS-PAGE gels (Bio-Rad Laboratories). In both cases, the migrated gels were transferred onto PVDF membranes (Millipore) and incubated overnight at 4 °C with corresponding primary antibodies (Table S1) in 50 mM Tris-HCl pH 8.0, 150 mM NaCl, 0.1 % Tween-20, 5 % milk. This was followed by incubation with horseradish peroxidase (HRP)-coupled secondary antibodies (1:10,000, Jackson ImmunoResearch) for 1 h at room temperature (RT) and revealed by chemiluminescence using Immobilon Crescendo or Forte Western HRP substrate (Millipore Merck).

For western blots against pMRLC, MRLC and actin, cells were lysed with 1x Laemmli (Bio- Rad Laboratories) with Benzonase nuclease and phosphatase inhibitors (Sigma-Aldrich p2850, Merck), and the lysates were migrated in 4–15 % gradient SDS-PAGE gels (Bio- Rad Laboratories). Gels were transferred onto nitrocellulose membranes (Whatman Protran), incubated overnight with primary antibodies in 50 mM Tris-HCl pH 8.0, 150 mM NaCl, 0.1 % Tween-20, 5 % BSA. fluorescently-coupled secondary antibodies (1:10,000, Jackson ImmunoResearch) were incubated for 30 min in Tris-HCl pH 8.0, 150 mM NaCl, 0.1 % Tween-20, 3 % milk at RT and revealed by LI-COR (Biosciences).

### Immunofluorescence and image acquisition

Midbody remnants were purified as described in (*43*) and fixed with pure methanol (#32213 Sigma-Aldrich) at -20 °C for 3 minutes. HeLa cells were grown on coverslips, and fixed with either 4 % paraformaldehyde (PFA, #15714 Sigma-Aldrich) for 20 min (Table S1) or pure methanol (#32213 Sigma-Aldrich) at -20 °C for 3 minutes (Table S1). Cells and midbody remnants were permeabilized with 0.1 % Triton X-100 and blocked with PBS containing 1% BSA for 1 h at RT, then incubated with the corresponding primary antibodies (Table S1) diluted in PBS containing 1% BSA for 1 h at RT. This was followed by an incubation with the secondary antibodies diluted in PBS containing 1% BSA for 1 h at RT. Cells were additionally stained with DAPI (0.5 mg/mL, Serva) for 10 minutes at RT. Cells and midbody remnants were finally mounted in Mowiol (Calbiochem). Image acquisition was done with an inverted Eclipse Nikon Ti-E microscope, using a 100x 1.4 NA PL-APO objective lens or a 60x 1.4 NA PL-APO VC objective lens, coupled to a CCD camera (Photometrics Coolsnap HQ) or Retiga R6 CCD Camera – Teledyne Photometrics and MetaMorph software (MDS), or using a CSU-X1 spinning disk confocal scanning unit (Yokogawa) coupled to a Prime 95S sCMOS Camera (Teledyne Photometrics) and CSU 100x objective. Images were then converted into 8- bit images using Fiji software (NIH) and mounted in Photoshop CS6.

### Osmotic shock experiments

Mild hyper-osmotic shocks were performed as in (*59*). Cells were seeded on glass coverslips the day before the experiment, and treated for 90 min either with isotonic cultured media (DMEM, 10 % FBS, 300 mOsm) or with DMEM, 10 % FBS media containing D-Glucose (#G5767 Sigma-Aldrich) to obtain a final osmotic pressure of 400 mOsm. Cells were then fixed and immunostained.

### Time-lapse microscopy

For time-lapse phase-contrast imaging (abscission times), cells were plated on glass bottom 12-well plates (MatTek) and put in an open chamber (Life Imaging) equilibrated in 5 % CO_2_ and maintained at 37 °C. Time-lapse sequences were recorded every 10 min for 48 h using an inverted Nikon Eclipse Ti-E microscope with a 20x 0.45 NA Plan Fluor ELWD controlled by Metamorph software (Universal Imaging).

For time-lapse fluorescent imaging, images were acquired using an inverted Nikon Eclipse Ti-E microscope equipped with a CSU-X1 spinning disk confocal scanning unit (Yokogawa) and with a EMCCD Camera (Evolve 512 Delta, Photometrics) or a Prime 95S sCMOS Camera (Teledyne Photometrics). Images were acquired with CSU 100x or CSU 60x 1.4 NA PL-APO VC objectives and MetaMorph software (MDS).

### Laser ablation

Laser ablation experiments were performed using a pulsed 355 nm ultraviolet (UV) laser (Roper Scientific) controlled by Metamorph and the iLas2 system (GATACA systems). The laser ablation system was connected to an inverted Nikon Eclipse Ti microscope confocal spinning disk equipped with a Yokagawa CSU-X1 spinning head, coupled to an EM-CCD camera (Evolve, Photometrics) and a 100x objective (Nikon S Fluor 100x 0.5-1.3 NA). The 355 nm UV laser is sufficient for membrane and microtubule cut, as described and demonstrated in [11]. To record the movement of the midbody following laser ablation, fast acquisition movies (20 ms time-lapse) were made using only transmitted light. The displacement of the midbody was then tracked using the software Icy (*67*). The midbody displacement after laser ablation follows a double exponential function as described in [11], and the retraction speed of the midbody was obtained from a linear regression of the first hundreds of ms after ablation.

### Cells synchronization

For EM and western blots for detecting pMRLC, MRLC and actin, a mitotic shake-off was performed to synchronize the cells in mitosis. Two days before the experiment, 2 million HeLa cells were seeded in T75 flasks. First, mitotic and dead cells were shaken-off and the remaining cells were further incubated for 2 h with fresh filtered media (DMEM GlutaMax supplemented with 10 % FBS and 1 % Penicillin/Streptomycin). Cells were again shaken-off, and the supernatant containing the mitotic cells was collected and centrifuged (900 xg for 3 min) in 15 mL tubes. The pellet of cells was then resuspended and added to collagen I (#A10483 Gibco) coated glass coverslips (50 μg/mL, for Electron Microscopy) and p6 well-plate (50 μg/mL, for Western blots), then incubated for 1 h and 2 h, respectively, at 5 % CO_2_ and 37 °C to allow them to exit mitosis and to progress into cytokinesis.

### Electron Microscopy

Glass coverslips with synchronized cytokinetic cells were chemically fixed for 2 h at RT in cacodylate buffer (0.1 M, pH 7.2) supplemented with 2.5 % (v/v) glutaraldehyde and 2 % (v/v) paraformaldehyde, then washed in cacodylate buffer (3 times), post-fixed with 2 % (w/v) osmium tetroxide supplemented with 1.5 % (w/v) potassium ferrocyanide (45 min for 4 °C), washed in water (3 times), dehydrated in ethanol (increasing concentration from 30 to 100 %), and finally embedded in Embed812/ Araldite (Epon) resin as described in (*68*). Microtomy of embedded cell monolayers was performed on a Reichert UltracutS ultramicrotome (Leica Microsystems) for 70 nm thin sections collected on formvar-coated cupper/ palladium grids, then contrasted with uranyl acetate and lead citrate. Electron micrographs acquired with a Transmission Electron Microscope (Tecnai Spirit G2; ThermoFischer Scientific, Eindhoven, The Netherlands) equipped with a 4k CCD camera (Quemesa, EMSIS, Muenster, Germany) using ITEM software (EMSIS).

### Image and Data Analyses

All images and videos were minimally processed to adjust the intensity levels, reduce image noise, and rotate the images with Fiji software. The kymographs (Fig. 4B) were obtained using an ROI (straight line) along the intercellular bridge at a fixed position over the entire movie, using Icy software. The scan lines were obtained using an ROI (straight line), with a width corresponding to the bridge or EP height, along the objects of interest, using Fiji. Pixel intensities values were extracted from the entry points defined with an ROI (square) for each channel of interest, using Fiji. Illustrations were created with BioRender.com.

### Statistical Analyses

All the plots and statistical tests were performed using GraphPad prism software. The presented values are displayed as mean ± SD (Standard Deviation) for at least three independent experiments (as indicated in the figure legends). The significance was calculated using unpaired, one-sided/two-sided t-tests, as indicated. For comparing the distribution of the abscission times, a nonparametric Kolmogorov–Smirnov test was used. In all statistical tests, a *P* > 0.05 was considered as non-significant. *P* values are indicated in the Figures.

## ACKNOWLEDGMENTS

We thank Matthieu Piel and the Echard Lab members for critical reading of the manuscript; Matthieu Piel, Aurélien Roux and Pierre Sens for helpful discussions throughout this project; and the Recombinant antibodies platform (TAb-IP, Institut Curie, Paris) for antibodies. We thank Adrien Presle and Nathalie Sassoon for the preparation of purified midbody remnants. We thank Audrey Salles from UTechS PBI, Institute Pasteur for discussions and advice in image acquisition; Maryse Moya-Nilges from UTechS PBI, Institute Pasteur for SEM training and help in image acquisition; Quentin Gaia Gianetto for making Fig. S1B; Pierre-Henri Commere from Cytometry and Biomarkers Utechs, Institut Pasteur for FACS sorting; Vincent Fraisier, the Cell and Tissue Imaging facility (PICT-IBiSA), and the Nikon Imaging Centre @ Institut Curie-CNRS for laser ablation experiments.

## Funding

Institut Pasteur (AE)

CNRS (AE)

ANR Cytosign, ANR SeptScort (AE)

French National Research Agency through the “Investments for the Future” program, France-BioImaging, ANR-10-INSB-04 (CD)

Cell and Tissue Imaging core facility (PICT-IBiSA), member of the France-BioImaging national research infrastructure, supported by the Labex Cell(n)Scale (ANR-10-LBX- 0038) part of the IDEX PSL (ANR-10-IDEX-0001-02 PSL) (CD)

Doctoral School Complexité du Vivant ED515, contrat n° 2828/2017 (VA)

La Ligue Contre le Cancer (4ème année de thèse) (VA)

Pasteur-Paris University (PPU) international PhD program (JB)

Fondation ARC pour la recherche sur le cancer (DOC20180507410) (JB)

## Author contributions

VA carried out and analyzed all the experiments except mentioned otherwise. CD performed, analyzed and interpreted the electron microscopy imaging experiments. JB obtained initial observations for Cavin1 localization. AJ taught laser ablation experiments to VA. NGR obtained initial data of the *Flemmingsome*. VA, CD, CL and AE designed the experiments. AE secured funding, analyzed and supervised the work. VA and AE wrote the manuscript with the help of CL and CD.

## Data and materials availability

All data are available in the main text or the supplementary materials.

---

**Figure S1.**
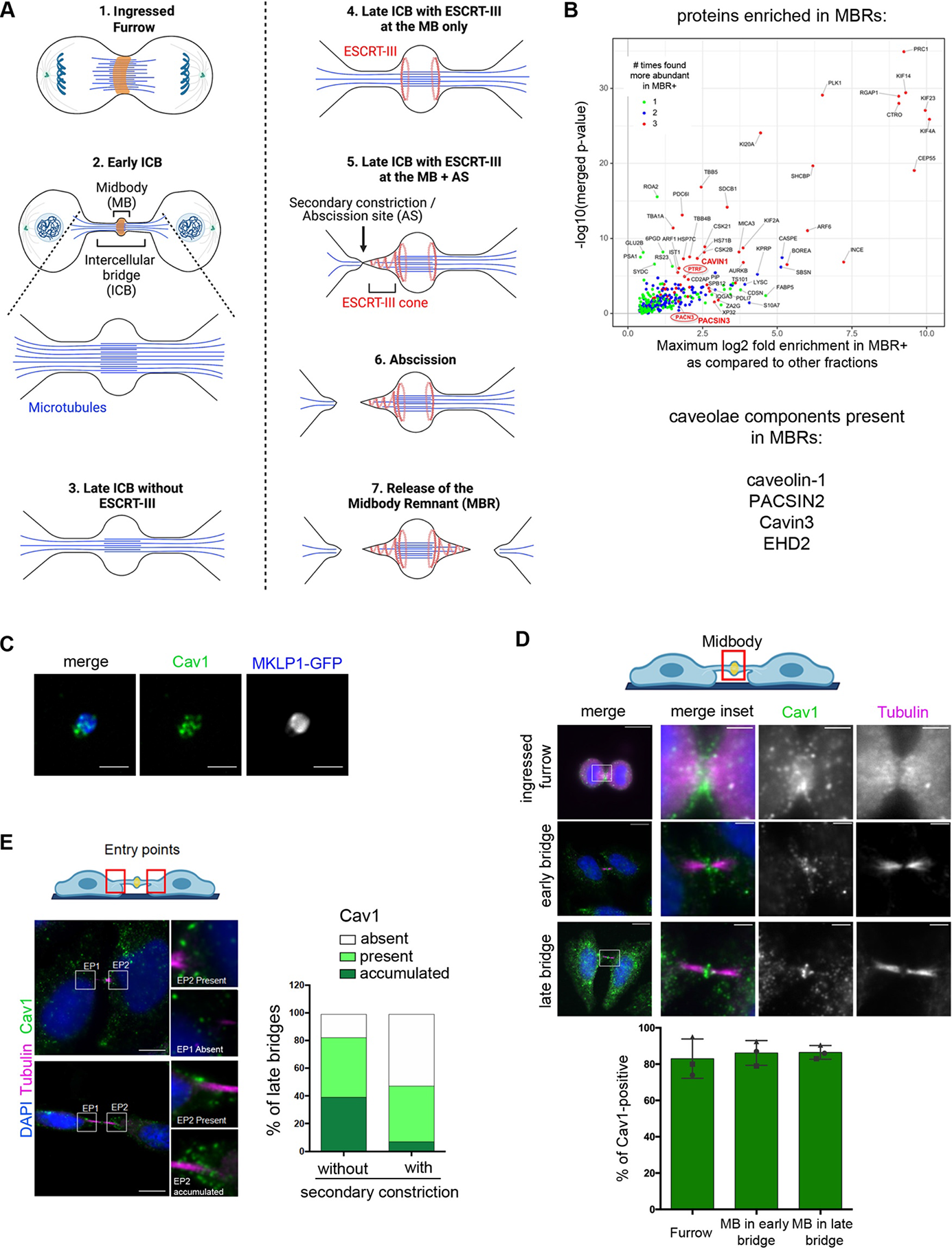
Cavin1 and Cav1 localization during cytokinesis. **(A)** Scheme depicting the successive phases of cytokinesis and the main definitions used in this paper. Note the different steps of ESCRT-III polymerization until abscission. A midbody remnant (MBR) is released after abscission on both sides of the midbody. **(B)** Merged volcano plot of the mass spectrometry analysis described and showing the *Enriched Flemmingsome* of HeLa cells. See details in (*43*). The caveolae components CAVIN1 and PACSIN3, which have been found enriched in MBRs are highlighted in red. Briefly, MBRs have been purified by flow cytometry from HeLa cells expressing the midbody marker MKLP1-GFP (MBR+ fraction). The plot shows the maximum log2(fold change) in *x*-axis measured between MBR+ purified fractions and the other fractions (either total cell lysates, MBRs enriched by centrifugation or GFP-negative fraction) and the corresponding –log10(merged p value) in *y*-axis. Color code: proteins significantly enriched in MBR+ when compared with 3 (red), 2 (blue), or 1 (green) of the other fractions. The other components of caveolae that are present but not enriched in the MBRs are listed below. **(C)** Purified midbody remnants from HeLa cells that stably express MKLP1-GFP were immunostained for endogenous Cav1. Scale bar, 2 µm. **(D)** *Upper panels:* Localization of endogenous Cav1, Tubulin, DAPI in HeLa cells at the indicated stage of cytokinesis. Scale bars, 10 µm (general views), 2 µm (inset). *Lower panel:* Percentage of the indicated structures positive for endogenous Cav1. Mean ± SD, n> 30 cells per condition. N = 3 independent experiments. **(E)** *Left panels:* Endogenous staining for Cav1, together with Tubulin and DAPI, in late bridges. Insets show different categories of Cav1 localization at EPs: accumulated, present or absent, as indicated. Scale bars, 10 µm. *Right panel:* Percentage of late bridges with/without secondary constriction where Cavin1 is accumulated, present or absent at EPs, as indicated. Mean ± SD, n> 40 cells per condition.

**Figure S2.**
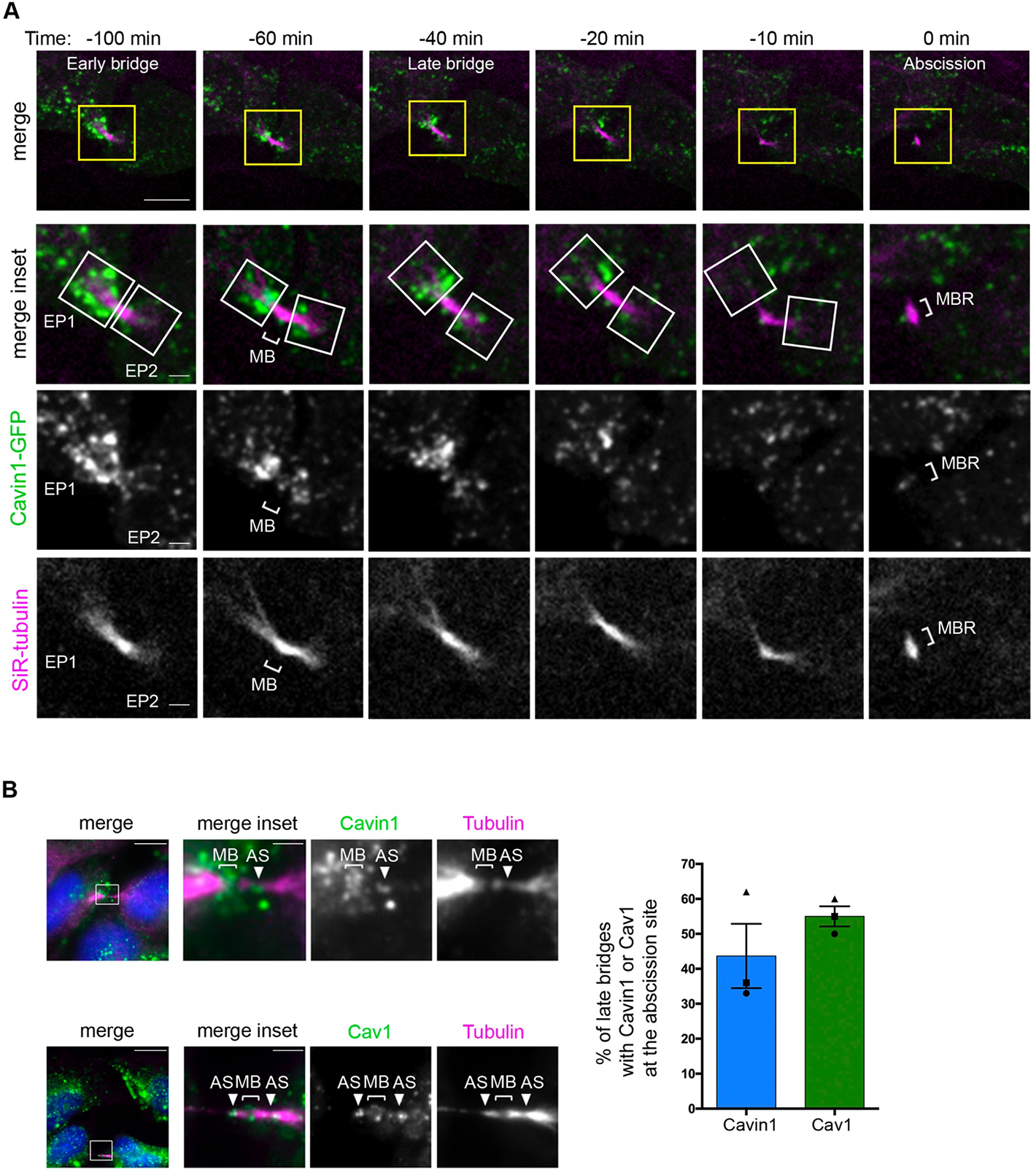
Cavin1 disappears from the EPs and the abscission site before abscission. **(A)** Snapshots of a spinning-disk confocal microscopy movie of a cell expressing Cavin1- GFP. SiR-tubulin was added for microtubule visualization. Regions shown in the insets correspond to the yellow boxes. Abscission occurred at Time 0 min. White squares delimit each EP. Note the progressive disappearance of Cavin1 from EPs as cells progress to abscission. Scale bars, 5 µm. **(B)***Upper left panels:* Endogenous staining for Cavin1, showing its localization at the midbody (bracket, MB) and at the abscission site (arrowhead, AS) denoted by the pinch in the Tubulin at this location. Scale bars, 5 µm. *Lower left panels:* Endogenous staining for Cav1, showing its localization at the midbody and at abscission sites. Scale bars, 5 µm. *Right panel:* Percentage of late bridges with Cavin1 or Cav1 localized at the abscission site. Mean ± SD, n> 30 cells per condition. N = 3 independent experiments.

**Figure S3.**
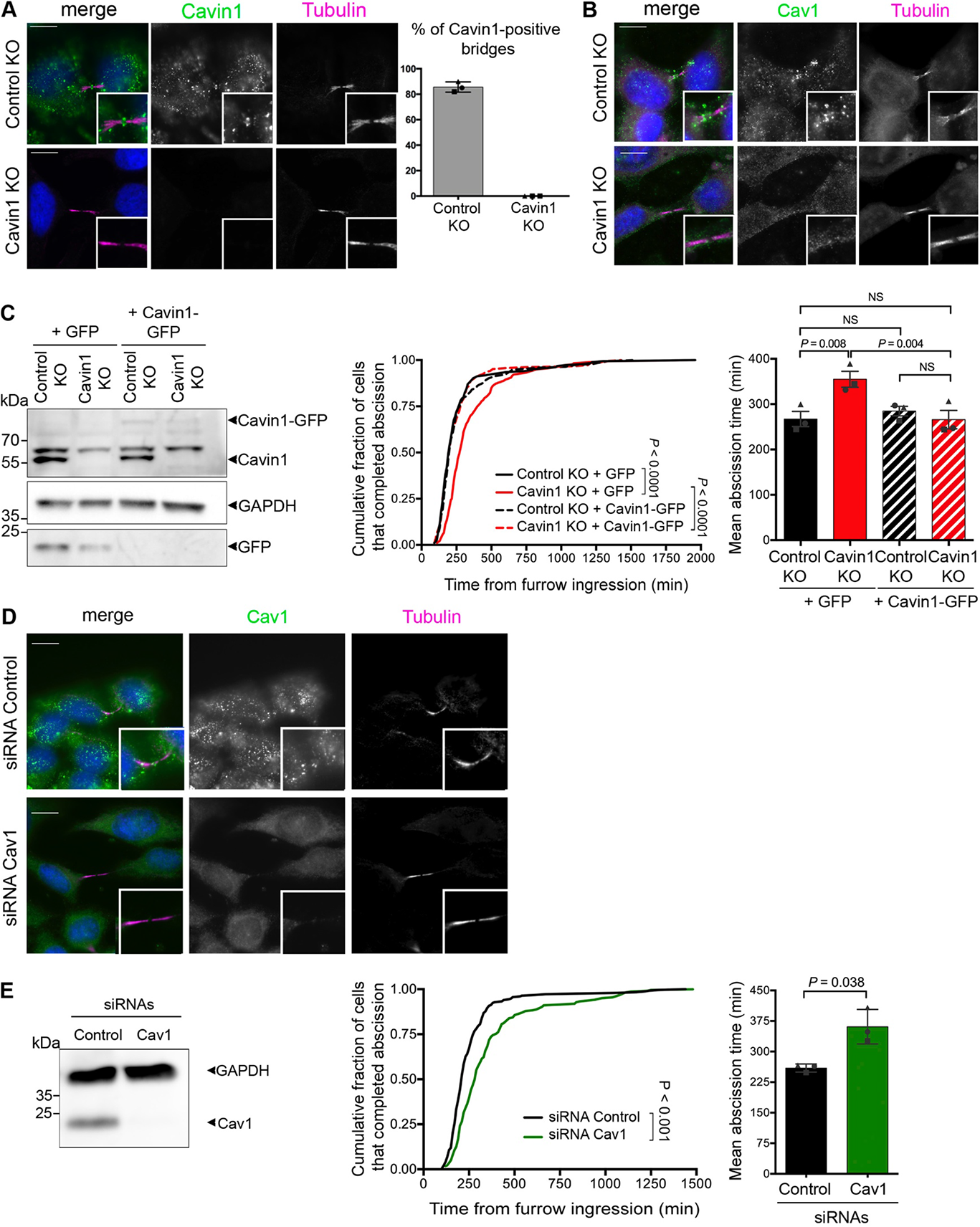
Cav1 depletion delays abscission. Specificity of the staining and rescue experiments. **(A)** *Left panels:* Endogenous staining for Cavin1, Tubulin and DAPI in Control KO and Cavin1 KO cells. Insets of the intercellular bridge are displayed. *Right panel:* Percentage of intercellular bridges that are positive for Cavin1 in Control KO and Cavin1 KO cells. N = 3 independent experiments, n> 20 cells per condition. Scale bars, 10 µm. **(B)** Endogenous staining for Cav1, Tubulin and DAPI in Control KO and Cavin1 KO cells. Insets of the intercellular bridge are displayed. Scale bars, 10 µm. **(C)** *Left panel:* Lysates from Control KO and Cavin1 KO cells transfected with either GFP or Cavin1-GFP, as indicated, were blotted for Cavin1 (upper part), GFP (lower part) and GAPDH (loading control). *Middle panel:* Cumulative distribution of the fraction of cells that completed abscission, measured by phase-contrast time lapse microscopy in Control KO and Cavin1 KO cells transfected with either GFP or Cavin1-GFP, as indicated. *P* values from nonparametric and distribution-free Kolmogorov–Smirnov [KS] tests. NS non-significant (*P* > 0.05). *Right panel:* Mean abscission times (min) ± SD in indicated cells, with n> 60 cells per condition. N = 3 independent experiments. Paired Student’s t tests. NS non-significant (*P* > 0.05). **(D)** Endogenous staining for Cav1, Tubulin and DAPI in Control and Cav1 siRNAs transfected cells. Scale bars, 10 µm. **(E)** *Left panel:* Lysates from HeLa cells transfected with Control or Cav1 siRNAs were blotted for Cav1 and GAPDH (loading control). *Middle panel:* Cumulative distribution of the fraction of cells that completed abscission, measured by phase-contrast time lapse microscopy in Control and Cav1 depleted cells. *P* values from nonparametric and distribution-free Kolmogorov–Smirnov [KS] tests. NS non-significant (*P* > 0.05). *Right panel:* Mean abscission times (min) ± SD in indicated cells, with n> 60 cells per condition. N = 3 independent experiments. Two-sided paired Student’s t tests. NS non- significant (*P* > 0.05).

**Figure S4.**
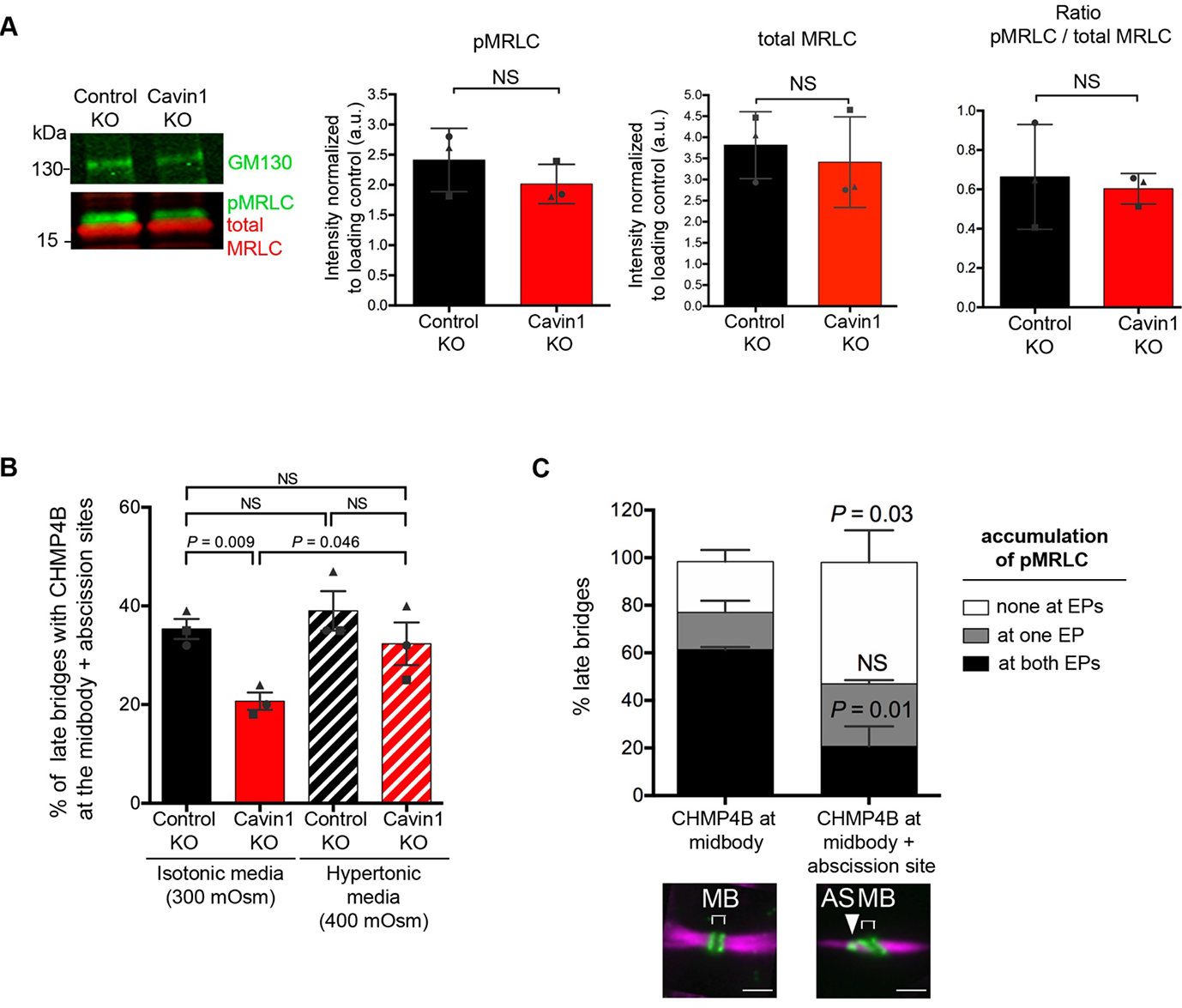
Global levels of activated myosin II and actin are unchanged in the Cavin1 KO cells. Hyperosmotic shock rescues ESCRT-III localization at the abscission site in Cavin1 KO cells. **(A)** *Left panel:* Lysates from Control KO and Cavin1 KO cells were blotted for pMRLC, total MRLC and GM130 (loading control). *Right panels:* quantification of the fluorescent westernblots presented above: pMRLC (normalized to GM130), total MRLC (normalized to GM130) and the ratio pMRLC / total MRLC. Mean ± SD, N = 3 independent experiments. Two-sided paired Student’s t tests. NS non-significant (*P* > 0.05). **(B)** Percentage of late bridges with CHMP4B localized both at the midbody and abscission site in Control KO and Cavin1 KO cells, either in isotonic media (300 mOsm) or hypertonic media (400 mOsm) for 90 min, as indicated. Mean ± SD, n> 34 bridges per condition. N = 3 independent experiments. Paired Student’s t tests. NS non-significant (*P* > 0.05). **(C)** Percentage of late bridges with pMRLC accumulation at one EP, two EPs or none at EPs, when CHMP4B is localized at the midbody only or both at the midbody and abscission site (see representative images, same image as in Fig. 3E), in wild type HeLa cells. Mean ± SD, n= 11-28 bridges per condition. N = 3 independent experiments. Paired Student’s t tests. NS non-significant (*P* > 0.05).

**Table S1.** Antibodies and fluorescent probes used in this study. MeOH: methanol fixation. PFA: paraformaldehyde fixation.

| **Protein** | **Company** | **Host** | **Name / clone** | **Dilution**<br>**WB** | **Dilution**<br>**IF** | **Fixation**<br>**IF** |
| --- | --- | --- | --- | --- | --- | --- |
| Cavin1 | Proteintech | rabbit | 18892-1-AP | 1/4000 | 1/2000 | PFA,<br>MeOH |
| CAV1 | SantaCruz | mouse | (7C8) sc-53564 |  | 1/100 | MeOH |
| CAV1 | Abcam | rabbit | ab2910 | 1/4000 |  |  |
| CHMP4B | Proteintech | rabbit | 13683-1-AP |  | 1/500 | MeOH |
| GAPDH | Proteintech | mouse | 60004-1-Ig<br>(1E6D9) | 1/40<br>000 |  |  |
| GM130 | Abcam | rabbit | EP892Y | 1/2000 |  |  |
| RMLC | Sigma | mouse | M4401 | 1/1000 |  |  |
| pMRLC | Cell Signaling | rabbit | antibody #3671 | 1/1000 | 1/100 | PFA |
| Actin | Chemicon, Merck |  | MAB1501 | 1/2000 |  |  |
| Tubulin | Institut Curie:<br>Therapeutic<br>recombinant<br>antibodies<br>platform | human | F2C-hFc2,<br>VHHD5-hFc1,<br>C3B9-hFc |  | 1/200 | PFA,<br>MeOH |
| Phalloïdin<br>488 | ThermoFisher<br>Scientific | N/A | N/A | N/A | 1/200 | PFA |
| Phalloïdin<br>642 | Cell Signaling<br>Technology | N/A | N/A | N/A | 1/200 | PFA |

## Supplementary Movies

**Movie S1**

HeLa cells transiently co-transfected with plasmids encoding Cavin1-GFP and Cav1-RFP incubated with SiR-tubulin were recorded by spinning-disk confocal microscopy every 10 min. Time 0 corresponds to the time of abscission. EP1: Entry Point 1; EP2: Entry Point 2. Scale bar= 5 μm

**Movie S2**

HeLa cells transiently transfected with a plasmid encoding Cavin1-GFP and incubated with SiR-tubulin were recorded by spinning-disk confocal microscopy every 10 min. Time 0 corresponds to the time of abscission. EP1: Entry Point 1; EP2: Entry Point 2. Scale bar= 5 μm

**Movie S3**

Control KO and Cavin1 KO HeLa cells were recorded by phase contrast time-lapse microscopy every 20 ms. Laser ablation was performed at 200 ms, as indicated. Note that the initial speed of retraction of the midbody is increased in the Cavin1 KO cell. Scale bar= 5 μm

